# Longitudinal multimodal neuroimaging after traumatic brain injury

**DOI:** 10.1101/2025.04.04.647315

**Authors:** Ana Radanovic, Keith W. Jamison, Yeona Kang, Sudhin A. Shah, Amy Kuceyeski

**Affiliations:** Department of Radiology, Weill Cornell Medicine, New York, 10065, NY, USA; Department of Computational Biology, Cornell University, Ithaca, 14850, NY, USA; Department of Mathematics, Howard University, Washington, 20059, DC, USA

**Keywords:** multimodal MRI, flumazenil PET, traumatic brain injury, longitudinal study

## Abstract

Traumatic brain injury is a major cause of long-term cognitive impairment, yet the mechanisms underlying recovery remain poorly understood. Neuroimaging methods such as diffusion MRI, functional MRI, and positron emission tomography (PET) provide insight into micro- and macro-scale changes post-TBI, but the relationships between regional cellular and functional alterations remain unclear. In this study, we conducted a longitudinal, multimodal neuroimaging analysis quantifying TBI-related pathologies in four biomarkers, namely flumazenil PET derived binding potential, dMRI-derived structural connectivity, and resting-state fMRI-derived functional connectivity and fractional amplitude of low-frequency fluctuations in individuals with mild-to-severe brain injury at the subacute (4–6 months post-injury) and chronic (1-year postinjury) stages. Brain injury related regional pathologies, and their changes over time, were correlated across the four biomarkers. Our results reveal complex, dynamic changes over time. We found that flumazenil-PET binding potential was significantly reduced in frontal and thalamic regions in brain injured subjects, consistent with neuronal loss, with partial recovery over time. Functional hyperconnectivity was observed in brain injured subjects initially but declined while remaining elevated compared to non-injured controls, whereas cortical structural hypoconnectivity persisted. Importantly, we observed that brain injury related alterations across MRI modalities became more strongly correlated with flumazenil-PET at the chronic stage. Regions with chronic reductions in flumazenil-PET binding also showed weaker structural node strength and lower amplitude of low frequency fluctuations, a relationship that was not found at the subacute stage. This observation could suggest a progressive convergence of structural and functional disruptions with neuronal loss over time. Additionally, regions with declining structural node strength also exhibited decreases in functional node strength, while these same regions showed increased amplitude of low frequency fluctuations over time. This pattern suggests that heightened intrinsic regional activity may serve as a compensatory mechanism in regions increasingly disconnected due to progressive axonal degradation. Altogether, these findings advance our understanding of how multimodal neuroimaging captures the evolving interplay between neuronal integrity, structural connectivity, and functional dynamics after brain injury. Clarifying these interrelationships could inform prognostic models and enhance knowledge of degenerative, compensatory, and recovery mechanisms in traumatic brain injury.

## 1 Introduction

Traumatic Brain Injury (TBI) is a leading cause of death and long-term disability world-wide. In those that survive, it is associated with multi-scale changes in brain anatomy and physiology. Acutely, there is neuronal death or dysfunction and axonal shearing at the cellular level.^1^ This is followed by continued neuronal death and dysfunction, remote degeneration of structural connectivity along white matter pathways,^1, 2^ and large scale shifts in functional co-activation patterns that have been associated with persisting symptomology or recovery of function. To date, it is not fully understood how microscale neurobiological changes relate to macroscale connectivity, knowledge which is crucial for gaining insights into injury and recovery mechanisms. Understanding mechanisms of TBI related change is paramount to developing effective therapeutics, of which there are none, and building more accurate prognoses, for which there are currently no accepted tools.

Approximately 70% of TBI survivors experience chronic cognitive impairments, particularly in attention, significantly impacting daily function.^3–5^ Prognostic models are generally lacking in sensitivity,^6, 7^ and there are currently no effective therapeutic interventions.^8^ The failure of most TBI clinical trials has been attributed to the insufficient knowledge of the pathophysiology of specific deficits which precludes development of targeted therapies, the inherent heterogeneity of TBI^9^ and the dearth of precise outcome markers^10^ to track spontaneous and intervention-driven recovery. Various neuroimaging tools such as magnetic resonance imaging (MRI; functional and diffusion) and positron emission tomography (PET) have been used to track and identify markers of functional and structural changes following TBI. Each modality provides a unique perspective on mechanisms of recovery after TBI, but, independently considered, they provide an incomplete picture.

PET is a powerful tool to investigate microscale changes post-TBI such as neurotransmitter availability, cellular metabolism, and molecular structural features. Here we use gamma -aminobutyric acid (GABA) receptor ligand ([11C]-flumazenil (FMZ) Flumazanil, (FMZ)) PET, a measure of GABA_A_ receptor availability that is thought to reflect neuronal loss, and has been used to study various psychiatric and neurological disorders.^11–15^ FMZ-PET has been shown to be sensitive to losses of neuronal integrity that correlate with cognitive impairment after TBI^13^ and in the chronic stage, TBI patients with persistent cognitive impairments exhibit low FMZ uptake in frontal cortices^12, 14^ and the thalamus.^12, 15^ These findings provide evidence of micro-scale changes from subacute to chronic post-TBI, where increases in binding might be an indication of repaired structure, or of increases in GABA_A_ activity in the remaining neurons.^16–20^ Although these observed changes could reflect mechanisms of recovery, it remains to be understood how these microscopic changes are reflected in macroscopic structural and functional differences measured via MRI.

Diffusion MRI (dMRI) is useful in studying white matter tracts which are commonly damaged by axonal shearing and/or lesions in TBI, while functional (fMRI) is useful in assessing brain co-activity pattern changes post-TBI. Although various measures can be derived from these modalities, commonly structural and functional connectomes (SC and FC, respectively) are extracted. SCs are matrices that reflect the strength of white matter connections between pairs of regions in some pre-defined atlas of gray matter structures, while FCs are matrices that reflect the co-activation of brain activity over time between pairs of gray matter regions in an atlas. Unimodal connectomic analysis after TBI has revealed links between SC losses and cognitive dysfunction^21–25^ and abnormal FC (either increases or decreases, depending on the study or region or network of interest) and persistent symptoms.^26–31^ Combined multimodal structural and functional connectomic analyses provide complementary information and, to date, have provided some insight into the connection between structural damage and functional outcomes.^28, 30, 32–35^ However, the heterogenenic and dynamic nature of TBI, as well as the effect of varied image acquisition, processing strategies, and inherent noise properties of f/dMRI, has led to conflicting results when mapping connectome changes to outcomes after TBI.

While resting state FC reveals co-activation changes between regions, fractional amplitude of low-frequency fluctuations (fALFF), provides complementary functional information regarding regional activity as it is a marker of the level of spontaneous neural activity.^36^ fALFF has been found to be increased in TBI patients,^37–40^ possibly reflecting neural dysfunction or excitotoxicity after TBI. Studies have linked increases in fALFF to increases in binding of fluorodeoxyglucose F-18 (FDG) and FMZ PET in controls,^41–43^ though the relationship is not well defined. fALFF’s relationship to connectomic measures have generally not been explored, especially in TBI, though fALFF and FC have shown positive correlations in order conditions such as autism^44^ None of these handful of studies examine relationships between fALFF and structural (SC), functional (FC), or FMZ-PET measures in TBI, let alone across recovery.

Intermodal relationships between MRI and PET markers (other than FMZ), such as FDG-PET (glucose metabolism), have demonstrated utility in identifying differential recovery trajectories in mild and moderate TBI subjects,^45^ elucidating chronic cognitive impairments in blast TBI,^46^ and predicting recovery in severe cases of TBI.^47^ Although multi-modal MRI and FMZ-PET analyses have provided insights into other conditions such as multiple sclerosis,^48^ its relevance to TBI recovery remains largely unexplored. In healthy non-injured adults, tri-modal studies (FMZ-PET, MRI, EEG) have linked increases in FMZ-PET binding to increases in resting fMRI measures, including fALFF, across networks like the executive control network.^42^ Also, utilization of multi-modal MRI and PET metrics have shown promise in distinguishing chronic TBI patients from healthy controls.^49^ These findings suggest joint FMZ-PET and MRI studies in TBI could reveal microscale (PET) and macroscale (MRI) mechanisms underlying recovery. Advancing knowledge of multi-scale mechanisms of TBI recovery trajectories requires the integration of microscale PET data with macroscale MRI features.

Here, we set out to relate diffusion and functional MRI metrics and FMZ-PET metrics longitudinally across the post-TBI recovery period to garner a more comprehensive understanding of multi-scale recovery mechanisms. We collected longitudinal PET and MRI in individuals with TBI at the subacute (4-6 months post-TBI) and chronic (1 year post-TBI) stages; imaging from non-injured controls was also collected. From each scan, we extracted multi-scale, regional biomarkers of brain anatomy and physiology at two levels: cellular, i.e. FMZ-PET binding potential (BP*_ND_*), and macroscale, i.e. diffusion MRI structural connectivity (SC), resting-state fMRI functional connectivity (FC) and fractional amplitude of low-frequency fluctuations (fALFF). We quantified TBI-related pathologies (compared to the non-injured controls) in the four biomarkers at both time points, as well as the change in these TBI-related pathologies over time. TBI-related regional pathologies, and their changes over time, were correlated across modalities. This paper is the first to compare cellular and macro-scale metrics of TBI-related pathologies to one another, and further, to investigate the evolution of these across-modality relationships over time.

## 2 Materials and methods

### 2.1 Study design

Multi-modal Flumazenil PET (FMZ-PET), diffusion (dMRI), and restingstate functional MRI (fMRI) were collected in individuals with TBI and non-injured healthy controls (HC). TBI subjects were imaged at 4-6 months after injury (mean = 150.5 days for MRI and 140 days for PET) and again 12 months after injury (average time between scanning sessions was 253 days for MRI and 297 days for PET). Healthy controls were imaged once for comparison to the TBI group. FMZ-PET data was collected from 7 TBI patients (4 female, aged 33–58, mean 48 years) and 19 HCs (7 female, aged 22–65, mean 44 years). DMRI and fMRI data were collected from 16 TBI patients (4 female, aged 19–73, mean 48 years) and 14 HCs (5 female, aged 23–86, mean 56 years); the 7 TBI PET subjects are a subset of the 16 TBI MRI subjects, and 9 HC PET subjects are a subset of the 14 MRI. Further demographic breakdown for subject numbers, sex, and age for each group and time point can be found in Supplementary Tables 1 and 2.

### 2.2 Participants and recruitment

TBI participants were recruited through inpatient rehabilitation units and trauma departments at large, urban academic medical centers. Control individuals were recruited through local advertisements. All study activities were approved by Weill Cornell Medicine’s Institutional Review Board.

All participants were required to meet the following criteria: (i) 18 years of age or above; (ii) English-speaking; (iii) capable of providing informed consent or a proxy/authorized agent available to provide informed consent; (iv) physically healthy and able to safely undergo PET and/or MR imaging; (v) not currently taking any psychoactive or benzodiazepine drugs; (vi) not currently taking any medication for attention-deficit/hyperactivity disorder; (vii) no history of schizophrenia, drug, or alcohol abuse; (viii) no history of epilepsy, stroke, dementia, or serious medical illness by self-report; and (iv) not pregnant (for female participants). Participants in the TBI group were recruited if they sustained a complicated mild (Glasgow Coma Scale 48 score of 13–15 with evidence of intracranial lesion as verified on acute neuroimaging) or moderate-severe TBI (Glasgow Coma Scale score *≤* 12) within the last 6 months.

### 2.3 PET acquisition and processing

We measured brain GABA_A_ receptor binding using PET imaging with the radioligand ethyl 8-fluoro-5,6-dihydro-5-[11C] methyl-6-oxo-4H-imidazo [1,5-a] [1,4] benzodiazepine-3carboxylate, or [^11^C]-FMZ. Binding potential (BP*_ND_*) to GABA_A_ receptors, a proxy measure for neuronal integrity, was extracted from the FMZ-PET images using PMOD software that implements the Logan graphical model with pons acting as reference region. See Kang et al., 2022^14^ for details on the PET acquisition and processing used in this study, though Kang et al., 2022^14^ used a simplified reference tissue model to estimate BP*_ND_*, rather than Logan graphical analysis.

### 2.4 MRI acquisition and preprocessing

A 3T Siemens Prisma scanner with a 32-channel head coil was used to collect anatomical, functional, and diffusion-weighted MRI images. The protocol was adapted from the Human Connectome Project Lifespan study.^50^ Anatomical images included 3D T1-weighted sagittal MPRAGE and T2-weighted SPACE (0.8mm isotropic voxels). Resting-state functional MRI was acquired with 2mm isotropic voxels, 72 axial oblique slices, 420 volumes, (TR/TE = 800/37ms, multi-band factor 8), with a total acquisition time of 11 min, 12 sec, divided between two scans with opposite phase encoding directions (AP = anterior-to-posterior, and PA = posterior-to-anterior). A matching pair of spin echo field maps with opposing phase encoding direction was collected for each resting state scan. Multi-shell diffusion-weighted MRI was acquired with 1.5mm isotropic voxels, 92 axial oblique slices (TR/TE = 3230/39.2ms, multiband factor 4), b=1500/3000, 92 directions per shell, acquired in both AP and PA phase-encoding directions for a total acquisition time of 22 mins 36 sec.

MRI data was preprocessed using the Human Connectome Project Minimal Preprocessing Pipeline.^51^ Anatomical images were inhomogeneity-corrected, anatomically segmented using FreeSurfer 6.0, and nonlinearly registered to the MNI152 template (6th generation). Functional MRI was motion-corrected, EPI distortion corrected using FSL’s “topup”,^52^ linearly coregistered to the anatomical image, and resampled to MNI152 space. Diffusion MRI was jointly corrected for motion, EPI distortion, and eddy current distortion using FSL’s “eddy” tool,^53^ before being linearly coregistered to the anatomical image. Regional analyses used FreeSurfer-generated, subject-specific 86-region atlases (68 cortical gyri from Desikan-Killiany, and 18 subcortical gray matter regions).^54, 55^

#### 2.4.1 Resting-state fMRI processing

Preprocessed functional MRI time series were denoised using custom scripts to identify outlier time points (motion derivative threshold 0.9 mm, global signal threshold 5*σ*), regress out motion and tissue-specific nuisance time series (24 motion time series^56^ and 10 eigenvectors derived from eroded white matter and CSF masks^57^), and temporally filter the final result (high-pass filter cutoff 0.008 Hz, using DCT projection). Outlier timepoints were excluded from nuisance regression and temporal filtering. Regional time series for each of the 86 gray matter areas was obtained from the denoised time series data. Pearson correlation between regional time series (excluding outlier timepoints) resulted in an 86×86 resting-state functional connectivity (FC) matrix for each AP and PA scan, which were then averaged together. Functional connectivity node strength for each region was computed as the sum of the positive elements in each row of the FC matrix (excluding negatives and diagonal).

fALFF is a metric that measures the relative contribution of the power of low frequency blood-oxygen-level-dependent (BOLD) signal fluctuations which are assumed to reflect the magnitude of neural activity.^36^ Spatial smoothing is first performed for every subject. Fast Fourier transform (FFT) was used to convert the signal to the frequency domain and the square root of the power spectrum was averaged across the 0.01–0.09 Hz domain. fALFF is the ratio of power in low-frequency band (0.0008-0.09Hz) to the power of the entire frequency range (0.008-0.625 Hz). fALFF was calculated at a voxel-wise level and then averaged to obtain regional fALFF.

#### 2.4.2 Diffusion MRI processing

Preprocessed diffusion MRI were further processed using MRtrix3,^58^ including bias correction, constrained spherical deconvolution (multi-shell, multi-tissue FOD estimation, lmax = 8^59^), and whole-brain probabilistic tractography (iFOD2^60^) using uniform white matter seeding (20 seeds per white matter voxel, resulting in approximately 5 million total streamlines per subject). Structural connectivity (SC) matrices were constructed by counting the number of streamlines that ended in each pair of gray matter regions, normalized by the total volume of each region pair. To mitigate the problem of false positive connections, region-pairs with zero streamlines in at least 50% of the uninjured control subjects were replaced with zeros throughout the dataset. Structural connectivity node strength for each region was computed as the sum of each row in the SC matrix (excluding the diagonal).

### 2.5 Statistics

Subacute (denoted as ses 1) and chronic (denoted as ses 2) analyses of each imaging metric were conducted via one-way analysis of covariance (ANCOVA) where t-statistics of coefficient’s group effects were examined as the primary effect of interest, with age and sex as covariates. All available healthy controls for a given modality were included in these ANCOVAs, which varied according to modality (see Figure 2, n = 7 TBI subjects had both MRI and PET). We included all available controls to increase power in determining TBI-related effects for a single modality, with an assumption that healthy controls are drawn from the same distribution within the population. A second set of ANCOVAs were performed using the subset of TBI subjects that had both MRI and PET data (n = 7) in order to compare MRI and PET measures. Each modality’s ANCOVA-based t-statistics for the group term (TBI vs control) were correlated via Spearman rank and p-values calculated using 1000 permutations. Longitudinal change within TBI subjects was analyzed using a paired t-test for only those with two time points (PET: n = 7; MRI : n = 16 TBI). All p-values reported are uncorrected for multiple comparisons. The analysis pipeline and methodology summary can be seen in Figure 1. Data were analyzed and visualized using custom scripts in using R (https://www.r-project.org) version 4.4.3 and Python.

**Fig. 1.**
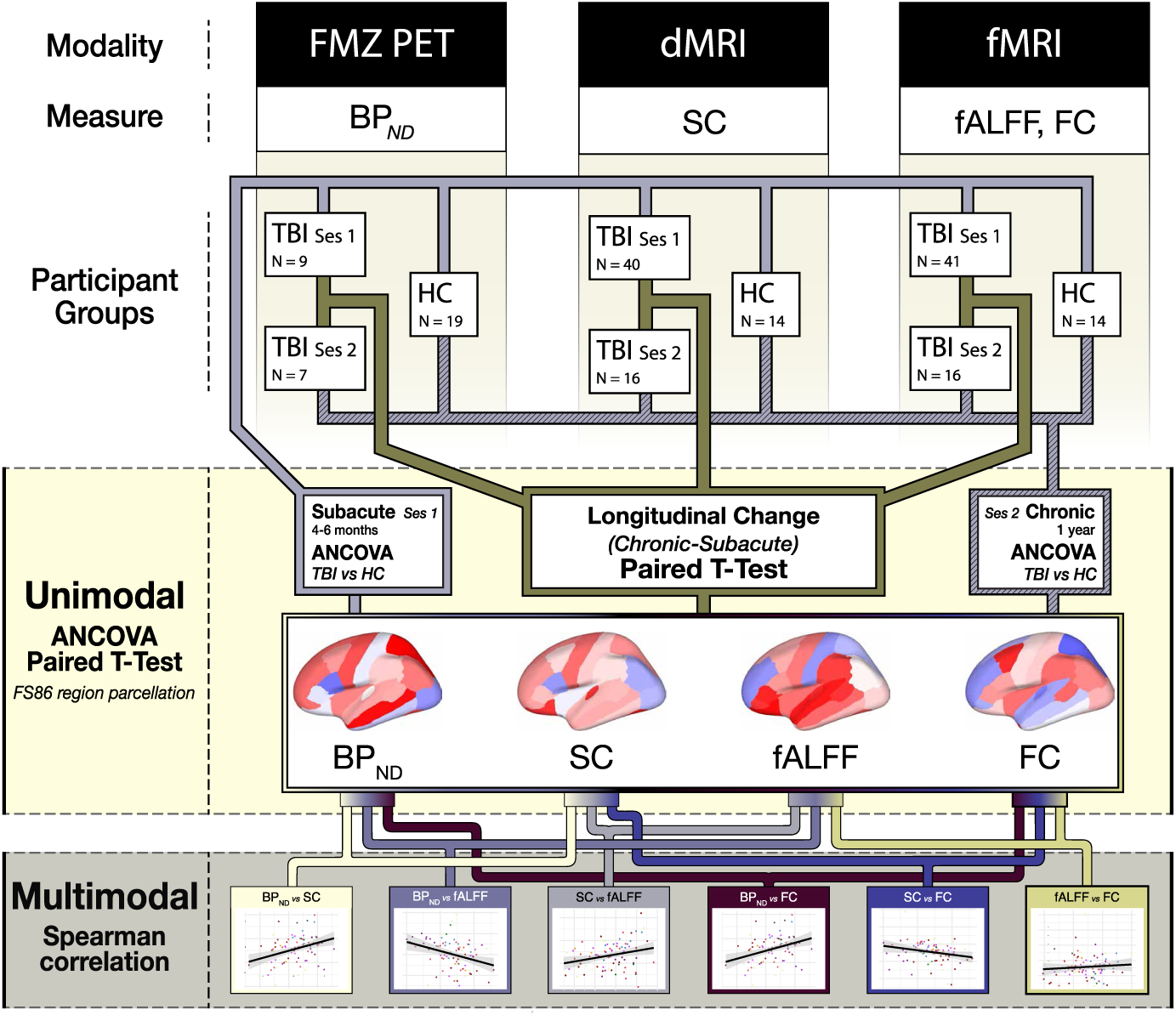
Analysis workflow for longitudinal multi-modal analysis. Between the 3 modalities (PET, dMRI, and fMRI) four markers per brain region were obtainednamely, BP*_ND_* from PET, SC node strength from dMRI, and fALFF and FC node strength from fMRI. These were extracted from all participant groups. TBI subjects have two sessions of recording 6-8 months apart with a single session from healthy controls. The number of subjects in each group and time point are denoted by N. ANCOVA was applied to uncover the effect of TBI on the various markers in the subacute (first) and chronic (second) session as compared to healthy, non-injured controls. A paired ttest between the 2 sessions was used to investigate changes over time in the TBI subjects. The t-statistics for the group coefficients (TBI vs controls) in ANCOVA and the t-statistics from the t-test from each of the four metrics are then Spearman correlated in a multimodal analysis.

**Fig. 2.**
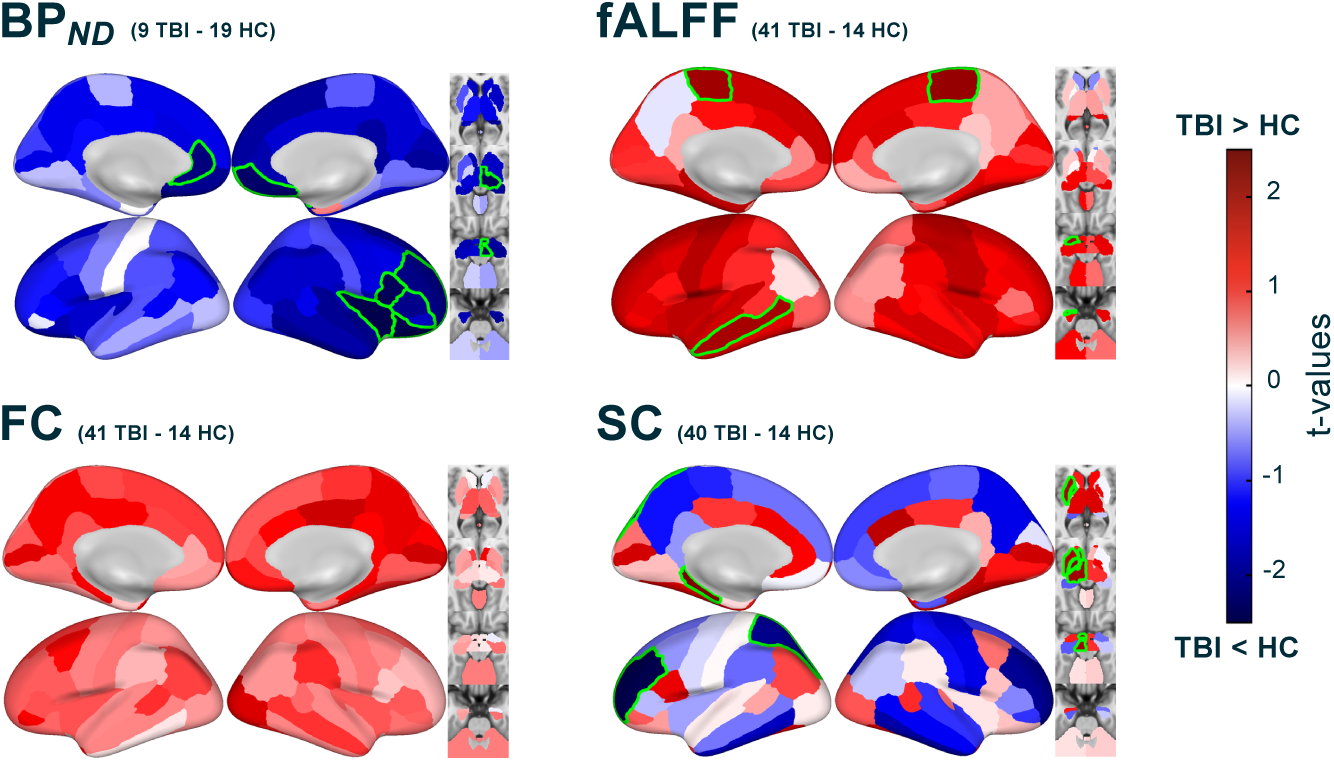
Group differences in [^11^C]FMZ tracer binding potential (BP*_ND_*), functional activity level (fALFF), functional connectivity (FC) node strength, and structural connectivity (SC) node strength in individuals with TBI compared to non-injured controls at the subacute timepoint. Subacute timepoint t-values of group coefficients from ANCOVA with age and sex as covariates. N varies by modality, as indicated in each subplot. Significant (uncorrected, p*<*0.05) values are highlighted in green.

## 3 Results

Group (TBI vs HC) coefficients’ t-statistics for each of the four biomarkers were plotted for each of the 86 regions for the subacute time point, see Figure 2. As reported before,^14^ [^11^C]FMZ - BP*_ND_* was generally lower in TBI subjects compared to controls, particularly in frontal and temporal regions as well as subcortical structures (p *<* 0.05). fALFF and FC exhibited the opposite pattern, in which TBI subjects showed generally higher values than healthy controls at the subacute timepoint. fALFF was significantly higher in sensorimotor and temporal regions while FC was most elevated in occipital and posterior cingulate (p *<* 0.05). Results for SC are mixed, with the cortex having generally lower structural connectivity (frontal, para-hippocampal, and parietal regions) reaching uncorrected significance (p *<* 0.05). A few regions in the left subcortex, including thamalus and basal ganglia, had trends for TBI being higher SC than HCs (p *<* 0.05).

For the longitudinal analysis, an ANCOVA was performed on a smaller subset of TBI subjects who had data collected at both time points (n = 7 PET, n = 16 MRI), see first and last rows Figure 3. The subacute results in the first row generally match the full dataset subacute results in Figure 2, with the exception of some regions having lower fALFF in TBI vs controls. The longitudinal change was analyzed looking only at the TBI subject’s change over time via a paired t-test, see the middle row of Figure 3. As reported previously,^14^ subjects with TBI had increases in cortical BP*_ND_* from subacute to followup; however, these increases were not large enough, generally, to return to the levels of non-injured controls, particularly in frontal regions. In the subcortex, although not significant, TBI subjects exhibit higher BP*_ND_* compared to controls in the cerebellum and left thalamus at the chronic stage. fALFF is initially higher in frontal regions and lower in parietal regions but over time, fALFF in frontal regions (particularly right hemisphere) and most subcortical regions decrease; there are concomittant increases in parietal regions. At the chronic timepoint, fALFF generally remains higher in TBI vs controls in the cortex but results are mixed in the subcortex.

**Fig. 3.**
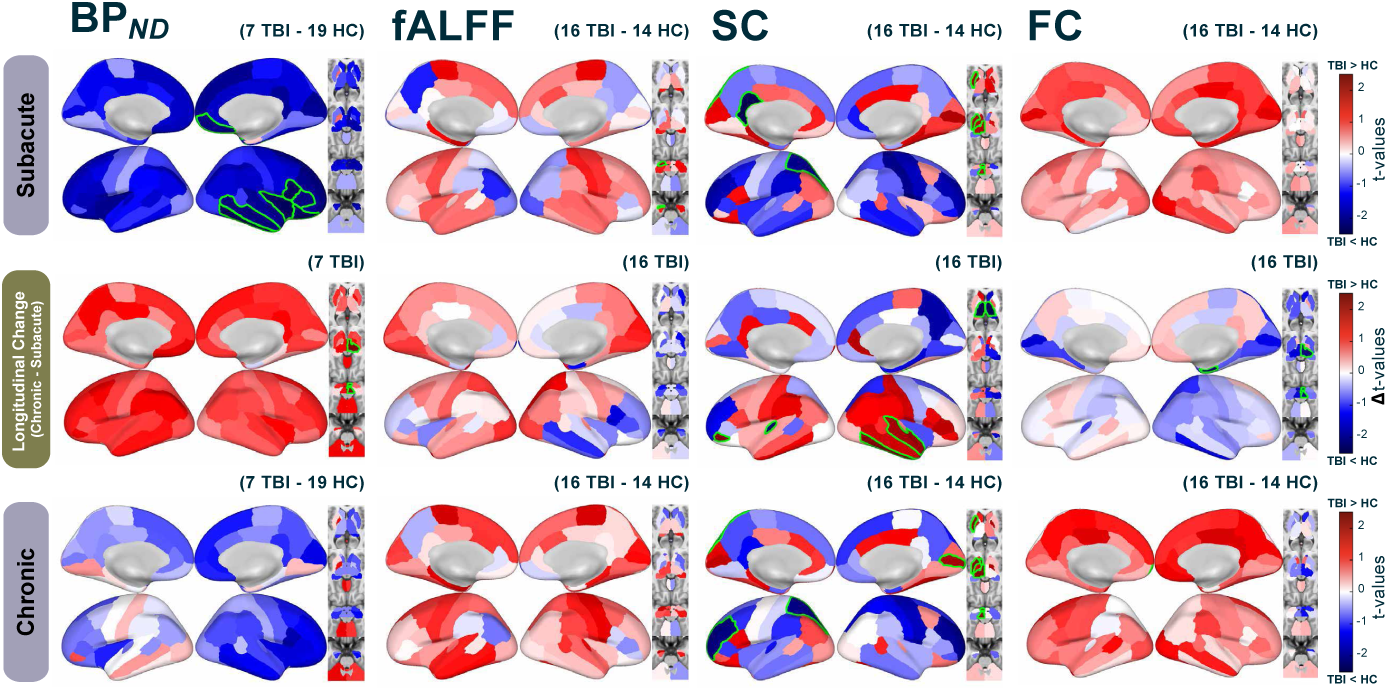
Longitudinal analysis of group differences (TBI vs controls) in regional FMZ tracer binding potential BP*_ND_*, functional activity strength (fALFF), structural connectivity node strength (SC), and functional connectivity node strength (FC). At baseline timepoint (Subacute - first row), longitudinal change (second row), and follow-up timepoint (Chronic - third row). Subacute and chronic timepoints are visualized via t-values of group coefficients from ANCOVA and longitudinal changes are visualized via t-values of paired t-test. N varies by modality, as indicated in each subplot. Regions with significant (p*<*0.05) values are outlined in green, uncorrected.

SC is generally lower in the subacute timepoint compared to controls, with more increases (trends toward significant changes in right temporal areas) than decreases over time, and generally still persistently lower SC at chronic stages, though less severe than in subacute stages. Patterns of TBI group differences in SC were very similar from subacute to chronic stages (see first and last row of Figure 3). Of note, in both the subacute and chronic timepoints, SC was significantly higher in the visual cortex, significantly so at chronic (p *<* 0.05, uncorrected). Thalamic regions that had initially higher SC at the subacute timepoint showed significant decreases in SC over time, while the putamen and other subcortical regions remained significantly higher compared to controls from subacute to chronic. FC was higher in TBI subjects compared to controls at the subacute timepoint, with reductions over time. Despite reductions in cortex FC over time, FC was still persistently larger than controls at the chronic timepoint. However, for some subcortical regions the reductions over time were enough to result in lower FC compared to controls at chronic.

Finally, we examined correlations of TBI-related differences in the four imaging biomarkers. As with the previous longitudinal analysis, only TBI subjects who had both sessions available were included (n = 7 PET, n = 16 MRI). Group ANCOVA coefficients’ (TBI vs HC) t-values from the multiple metrics at each time point were correlated via Spearman rank correlations; paired tstatistics from the change in one metric to the change in the other metric were also Spearman rank correlated, see Figure 4. In the first column, BP*_ND_* and fALFF are positively correlated in the chronic time point in the cortex (r = 0.28, p = 0.02), such that regions with larger TBI-related decreases in binding potential also had fALFF values that were not as elevated in TBI compared to other regions. Trending but not significant at the subacute timepoint, this relationship is opposite to what is seen in the subacute timepoint, where regions with more TBI-related decreases in BP*_ND_* also had more elevated fALFF. In the second column, change in BP*_ND_* and change in FC in cortical regions are positively correlated (r = 0.23, p = 0.03), indicating regions with larger increases in FMZ-binding also had larger increases in FC. In the third column, BP*_ND_* and SC are positively correlated at chronic timepoint (r = 0.23, p = 0.03) and trended toward significance at the subacute timepoint, such that regions with larger TBI-related decreases in binding also had more SC damage. In the fourth column, fALFF and FC are not correlated at subacute or chronic timepoints, however, the longitudinal changes in fALFF and FC were negatively correlated (r = -0.26, p = 0.04) in cortical regions such that regions with more increases in fALFF over time also had larger decreases in FC. Although not significantly negatively correlated at the subacute and chronic timepoints, the longitudinal change in fALFF and SC were negatively correlated in the cortex (r = -0.33, p=0.01) such that regions with decreasing SC had increasing fALFF. In the final column, FC and SC are positively correlated in cortical regions at both subacute (r = 0.32, p = 0.01) and chronic timepoints (r = 0.26, p = 0.04), with longitudinal change within regions trending toward a positive correlation meaning regions with larger TBI-related decreases in SC also had less TBI-related elevations in FC.

**Fig. 4.**
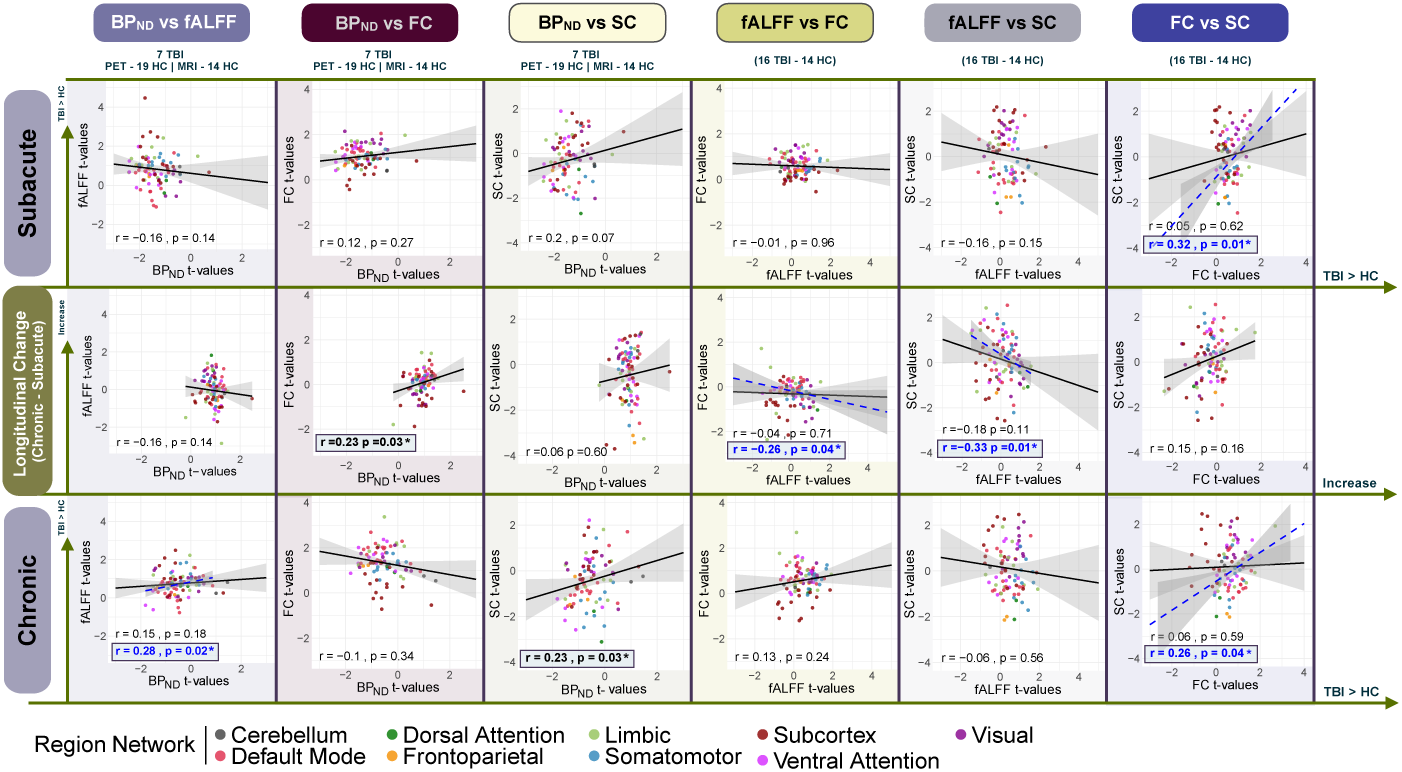
Inter-modality comparison of TBI-related changes in imaging metrics. Spearman correlations of the t-values from ANCOVA group coefficients (TBI vs control) between FMZ tracer BP*_ND_*, fALFF, FC, and SC at baseline (Subacute - first row) and follow-up (Chronic - third row). T-statistics indicating change vs change for the two modalities are in the middle row. Correlations are calculated for all brain regions or, where indicated, cortical regions only (in blue). Cortex-only correlations (dashed blue line) are only shown for comparisons where it differs substantially from the whole-brain correlation (solid black line). Correlation strength and p-value indicated in bottom corner, asterisk and bold indicate significance (uncorrected, p*<*0.05). N varies by modality, indicated under titles of paired modality columns. Each region is colored by the functional network to which it belongs (see legend).

We provide additional details on the main results in the Supplementary Information (SI), including robustness analyses and global correlations between imaging metrics, cognitive scores, and outcome measures. First, ANCOVA *η*^2^ results for group effects of t-values plotted in Figures 2 and 3 are shown in Supplementary Figure 3. Next, we performed a multi-modal correlation analysis on a larger cohort, including all available subjects at the subacute timepoint, to increase statistical power. The *t* -values from Figure 2 were correlated across modalities (see Supplementary Figure 1. Consistent with findings from the smaller cohort (Figure 4), TBI-related differences in FC and SC node strength were positively correlated in the cortex. Trends toward positive correlations between TBI-related BP*_ND_* and SC node strength were observed in both cohorts, though neither reached significance. These results suggest consistency in the observed relationships across different sample sizes. Additionally, we examined global average imaging metrics in relation to cognitive and standard TBI outcome measures. Overall, no significant associations were found, except for a few uncorrected relationships where increased fALFF and BP*_ND_* were linked to improvements in GOSE scores over time. However, the relationships between individual global metrics and various outcomes at subacute and chronic timepoints were inconsistent and did not reflect a clear pattern across different outcome measures (see Supplementary Figure 4).

## 4 Discussion

Here we use multi-modal PET and MRI imaging to understand multi-scale structural and functional changes during recovery from TBI. While our unimodal findings align with prior research, we find that the across-modality relationships of TBI-related differences allow deeper insight into the complex and dynamic recovery process. Overall, the across modality (PET-MRI) relationships were stronger in the chronic stage. We found that regions with chronic, persistent decreases in BP*_ND_* due to TBI also had weaker SC node strength and less elevated neuronal activity measures, a relationship that wasn’t found at the subacute time point, suggesting a convergence of structural and functional disruptions driven by neuronal loss over time. We also found that regions with TBI-related decreases in SC node strength also had TBI-related decreases in FC node strength, and furthermore, regions with decreasing SC and FC node strength over time also had increases in a measure of neural activity (fALFF) over time. The latter potentially indicating neural hyperactivity or a compensatory mechanism in regions that are increasingly disconnected from the rest of the brain.

### 4.1 Longitudinal unimodal changes during recovery

As shown before by our group and others,^14, 15^ we found lower BP*_ND_* in TBI subjects compared to controls, with an increase over time to levels that were still generally lower than controls, particularly in frontal regions and the thalamus. It has also been found that lower binding in thalamic regions is associated with worse outcome after TBI.^15^ Importantly, some regions that had significantly lower binding such as the anterior cingulate cortex (ACC) are heavily implicated in attention and emotional processing, which are common afflictions post-TBI.^4, 61–64^

Functional connectivity analyses in resting state fMRI after TBI have been conflicting.^65–67^ Typically analyzed via seed or ICA based methods, studies report resting state hyper-connectivity^27, 32, 68^ in the very acute stages (within 2 weeks of injury) between the thalamus and the pre-frontal cortex,^27, 28^ and the DMN and the frontal cortex.^29^ Other research suggests that hypoconnectivity within the DMN and hyperconnectivity with the DMN and other regions at rest in acute and chronic stages, that can predict persistent cognitive symptoms.^30, 31^ At 6 months after injury, higher FC is associated with lower cognitive impairment.^30^ Generally, however, higher FC is found after TBI which aligns with our findings at both time points - at least in the cortex (see Figures 2 and 3. It is thought that the brain attempts to compensate for structural and functional disruptions by recruiting resources from other areas, i.e. functional compensation.^31, 32, 37, 68, 69^ Despite the generally higher FC in the cortex at both time points, subcortical structures had higher FC in the subacute stages but significant decreases over time, with some regions having lower FC compared to controls at the chronic time point. This could potentially reflect increased vulnerability of the connections between subcortical and cortical regions^15^ that worsen over time and result in their decoupling. However, aside from heterogeneity in TBI severity and localization, different choices in fMRI preprocessing (i.e. global signal regression; GSR) and calculation of FC (seed-based, atlas-based, correlation type, etc) complicates cross-study comparisons. We found an overall increase in FC in subacute TBI which gradually decreases over time but remains elevated compared to controls, except in subcortical areas where FC decreases over time to levels below controls.

Increasing fALFF over time has been correlated with improved symptom scores from 0 to 3 months after TBI,^70^ complementary to a finding of lower fALFF in the thalamus, frontal, and parietal regions at *<* 25 days after injury in subjects with post-concussive symptoms.^71^ Our analysis showed subjects had generally higher fALFF across cortical regions, except for occipital regions, some subcortical areas, and cerebellum at the subacute stage. Individually, higher fALFF and FC values are thought to possibly reflect compensatory mechanisms for structural damage^31, 32, 37, 68, 69, 72^ and reduction in these values over time may signal an exhaustion of these compensatory mechanisms, or a relief due to recovery. Also, decreases in fALFF in some regions over time, as was found in right frontal and temporal areas, could possibly reflect the loss of neurons due to damage.^73^ Similarly, decreasing fALFF over time in the thalamus could reflect progressive injury, which has been found after TBI.^15^ Large, comprehensive longitudinal studies with (1 year +) multiple measured timepoints may be able to better capture the dynamics of these mechanisms. SC node strength was generally lower in TBI compared to controls at both subacute and chronic timepoints, see Figures 2,3, in line with the idea that injury induces not only white matter tract damage but also progression of continued degeneration along axonal tracts.^74^ SC analysis has often found hypoconnectivity due to damage,^22^ particularly in temporal and frontal white matter tracts, that has been associated with cognitive dysfunction.^75^ SC hyperconnectivity has also been identified in “central hub areas” in very acute timepoints, which could either reflect the effects of acute edema^76^ or potentially effects of functional reorganization with information processing to higher hubs, akin to rich club networks.^32^ We found that some regions in the visual cortex and left putamen exhibited higher SC over time in TBI compared to controls, with regions in the right (and to a lesser extent left) temporal had increases in SC over time, which could be reflective of either repair to white matter tracts or changes in response to the widespread functional upregulation. The thalamus showed trends toward initially increased SC with significant decreases in SC over time, in line with findings that the thalamus, although not necessarily impacted directly by the initial insult, will show progressive neuronal and structural loss through TBI recovery.^15^ SC differences are also difficult to tease apart as well because of the heterogeneity in TBI etiology and spatial patterns of axonal injury across subjects.

### 4.2 Multi-scale PET and MRI relationships

Very few studies have related FMZ-PET to (f)MRI metrics. One prior research finding in controls using simultaneous PET-MR-EEG imaging found network level correlations across individuals between fALFF and BP*_ND_*,^42^ wherein higher fALFF was related to higher BP*_ND_*. Other research found a positive correlation between BP*_ND_* and increases in FC in the visual cortex between changes in eyes open and eyes closed conditions at rest.^77^ Here, we identified a positive correlation between TBI-related pathology in fALFF and BP*_ND_* at the chronic timepoint, opposite to the relationship at the subacute timepoint. This somewhat paradoxical finding could reveal neural dysfunction that results in hyperactivity at the subacute point, which, alongside chronic neurodegeneration,^1^ exhausts at the chronic stage more dramatically for those neurons with dysfunction or death (decreased BP*_ND_*) resulting in lower levels of neuronal activity reflected by lower fALFF. In other work, a related measure called ALFF was shown to be positively correlated with FMZ-PET and glucose metabolism (rmGLU)-PET in healthy controls and some subjects with temporal lobe epilepsy (TLE), but not in all subjects. As ALFF was found to be more highly correlated with rmGLU than FMZ-PET BP*_ND_*, decoupling of ALFF and FMZ-PET may be due to impaired neurovascular coupling^78^ which could be a factor more salient subacutely in our TBI population.^79^

FMZ-PET binding has been shown to correlate with MRI-based atrophy in epilepsy^80^ and with T2-weighted lesion load in multiple sclerosis,^48^ and to possibly capture neuronal loss not visible on anatomical MRI.^13^ However, the relationship between FMZ-PET binding and dMRI-based SC has been explored in only a handful of studies. One such prior study found that individuals with lower thalamic FMZ-PET binding had more lesions in the white matter connecting to the thalamus (i.e. weaker SC).^15^ We found an alignment of this previous work in that at both timepoints, BP*_ND_* and SC were positively correlated, i.e. regions with more TBI-related decreases in FMZ-BP*_ND_* also had more decreases in SC node strength (see Figure 4). This relationship only reached significance at the chronic time point, which could be due to continued neuronal degeneration after TBI that resulted in a strengthening of the relationship between the two metrics. TBI effects on MRI and PET measures are generally more correlated at the chronic timepoint. Our conjecture is that there is perhaps TBI-related neuronal loss occurring over time which is reflected in persistently lower BP*_ND_*, longitudinal decreases but still higher-than-normal fALFF and FC, and weakening SC node strength.

### 4.3 MRI-based structure-function relationships in TBI

FMRI provides information on brain activity (e.g. fALFF) and co-activity patterns (FC), while diffusion MRI provides estimates of the amount of structural (white matter) connections between regions (SC). The relationship between fALFF and measures of connectivity are not well defined, however the relationship between SC and FC has been explored widely across health and disease/disorder.^33, 68, 81–89^ We found that longitudinal change in fALFF and FC+SC were negatively correlated in the cortex, wherein regions with larger decreases in FC and/or SC over time also tended to have increases in fALFF. We conjecture that increasingly functionally and structurally disconnected regions had increasing (decoupled) neuronal activity, either as an effect of broad axonal degradation or less regulation from outside inputs.

ALFF and FC have been positively correlated in autism^44^ and in controls and individuals with mild cognitive impairment.^90, 91^ However, in both studies, this was limited to certain regions; most regions did not have such correlations between the fMRI metrics. Prior research has found a positive correlation in controls between SC and relative low frequency power,^92^ a related measure to fALFF, a correlation we also found to a degree in HC and TBI subjects when we looked at regional values (not the TBI related effects on the metrics, see Supplementary Figure 2). As for the relationship between SC and FC, a longitudinal study found that at the acute timepoint (within 7 days of injury), FC and SC were negatively related.^32^ Similarly, another study at the chronic time-point showed decreased SC in the cingulum was correlated with increased FC within the frontal node of the DMN.^73^ Our results are somewhat contradictory to these prior findings, but our analysis is inherently different, as we correlate regional node strength changes in structure and function due to TBI.

### 4.4 Limitations and Future work

There are some important limitations to this study, the first and foremost being the small sample size, particularly for PET imaging (only 7 TBI subjects had both subacute and chronic PET and MRI data). PET is expensive and difficult to collect and analyze, and involves intravenous injection of a radiotracer so it poses more risk to individuals. Both (f)MRI and PET are modalities suffer from low SNR, particularly in the subcortex, making it more difficult to obtain reliable measures in those regions. Secondly, as with many TBI studies, the individuals included in our cohort have wide heterogeneity in injury location, etiology and severity. Although there is a range of injury severity in our cohort, mild TBI was more common in our study 1 so any of our findings here may not reflect potentially different mechanisms in severe TBI. Thirdly, distributions of GABA-A receptors (which is where FMZ-PET binds) varies across the brain and thus our measure of BP*_ND_* may be impacted by this heterogeneity.^42^ Our main longitudinal analysis also did not have one important factor, namely, a chronic time point for the non-injured controls. Including such information would allow isolation of TBI-related changes over time compared to what could be test-retest measurement noise or natural variability in individuals. Also, other work including graph theoretical measures of SC and FC and their inter-relationship could be used to reveal differing topological changes due to TBI.^34, 93, 94^ Finally, future work should look at how relationships between metrics, perhaps within specific networks, are related to various measures of outcome.

## 5 Conclusion

Here, we use longitudinal multi-modal MRI and FMZ-PET imaging in TBI to examine cross-modal relationships during recovery after TBI. We found that individually, patterns of FC, fALFF, SC, and BP*_ND_* generally agreed with previous unimodal studies. Multimodally, however, the relationships between the metrics revealing TBI effects over recovery proved to be dynamic and complex, with an increasing convergence of MRI and PET markers over time. FC and fALFF, as thought to be reflective of possible compensatory mechanisms, were found to be increased subacutely and decreasing over time, resulting in better alignment with structural and neuronal damage in the chronic stage. Taken together, this longitudinal multi-modal study is a first step into understanding not only dynamics in recovery, but how multi-scale structure and function relate across recovery after TBI.

## A Data availability

Derived data and code can be made available upon reasonable request.

## B Acknowledgments

We gratefully acknowledge the research support of Ryan J. Lowder, Abigail Patchall, Karen Wen and Drs. Gary Dorfman, Doug Ballon and Susan Gauthier. We also acknowledge Franck Porteous for his support in visualization.

## C Competing Interests

The authors report no competing interests.

## D Funding

This work was funded by NIH grants R01 NS102646 (AK and SS), R01 NS134646-01A1 (AK), RF1 MH123232 (AK), and R01 AG077576 (SS). This work also used REDCap, which is supported by the Weill Cornell Clinical and Translational Science Center (CTSC) with funding from NIH grant number UL1TR024996.

## E Supplementary information

### E.1 Multimodal correlations large group

We repeated the multi-modal correlations of TBI-related pathology at the subacute timepoint using the larger dataset of all individuals with subacute fMRI and dMRI data. This was only done with dMRI and fMRI because subjects with both PET and MRI were subjects who also had both sessions of data, therefore, there are no new subjects to analyze in this set. T-values of the coefficient’s effects from the regional subacute unimodal ANCOVA analysis in Figure 2 were correlated across the neuroimaging markers via Spearman rank correlation with permutation-based p-values. Correlations were calculated for all 86 cortical, subcortical and cerebellar regions (solid black line) and for 68 cortical regions only (dashed blue line) to isolate correlations in the cortex from whole brain correlations. Results are depicted in Supplementary Figure 1. A similar relationship is observed between FC and SC, such that the TBI-related pathology in FC and SC was positively correlated in the cortex (r = 0.32, p = 0.01), such that regions with lower FC in TBI compared to HC were also found to have lower SC compared to HC. This was in the same direction as what was found with the smaller cohort in the main text. Both the smaller cohort and the larger cohort had trends for negative correlations between TBI-related fALFF changes and TBI-related SC changes although neither were significant.

### E.2 LFO and SC node strength

Previous work has identified correlations between fMRI-derived low frequency oscillations (LFO) and the structural connectivity strength of the region in non-injured controls.^92, 95^ Here, using the non-injured controls (n = 14) and TBI subjects (n = 40) separately, we correlated each regions’ SC node strength with the same regions’ fALFF values via spearman correlation, and found a significant positive correlation (TBI: r = 0.23, p = 3.72 * 10^-44^, HC : r = 0.29, p = 7.05 * 10^-25^) which agreed with those previous works’ findings (see Supplementary Figure 2). The p-values reported are permutation based.

**Supplementary Figure 1.**
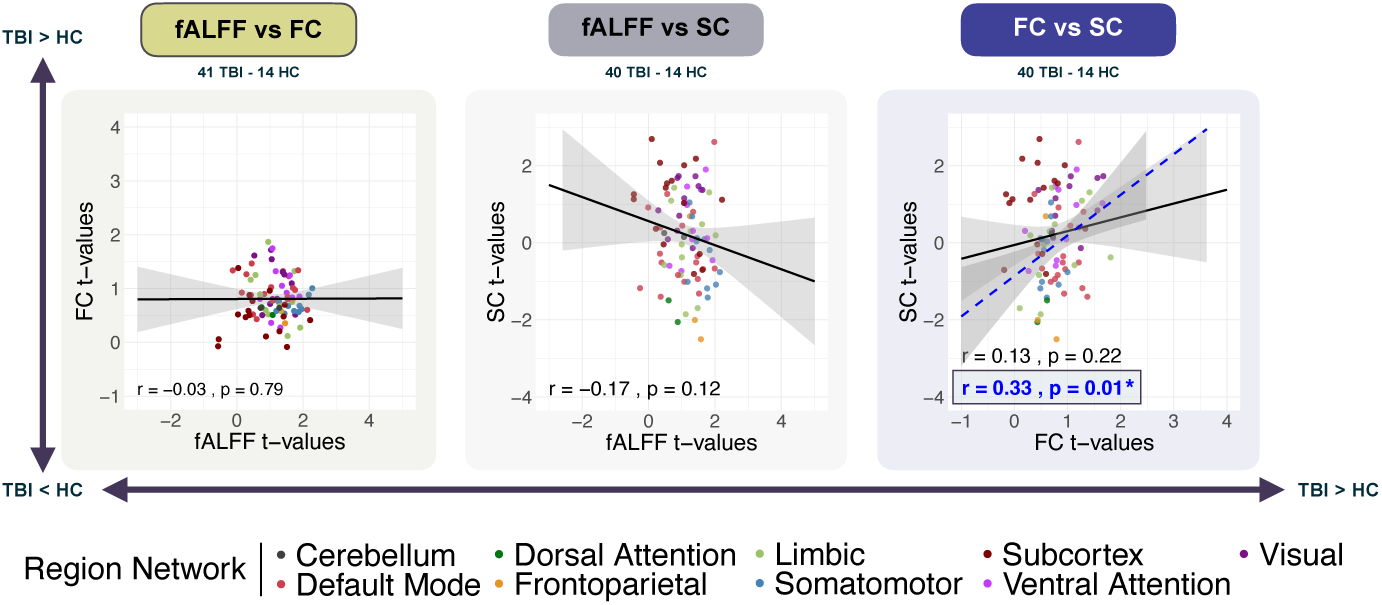
Inter-modality correlations of the t-values between functional activity strength (fALFF), functional connectivity (FC), and structural connectivity (SC) in all individuals with TBI at the subacute timepoint compared to controls. Subacute timepoint inter-modality correlations of the t-values of TBI group effects from ANCOVA analysis via Spearman correlation with permutation-based p-values. Correlation of group effects for whole brain or, where indicated, cortex only (in blue). Cortex-only correlation only shown for comparisons where it differs substantially from the whole-brain correlation. Correlation strength and significance highlighted in bold with a box and asterisks for comparisons with p*<*0.05 (uncorrected). N varies by modality, as indicated in each subplot.

**Supplementary Table 1.**
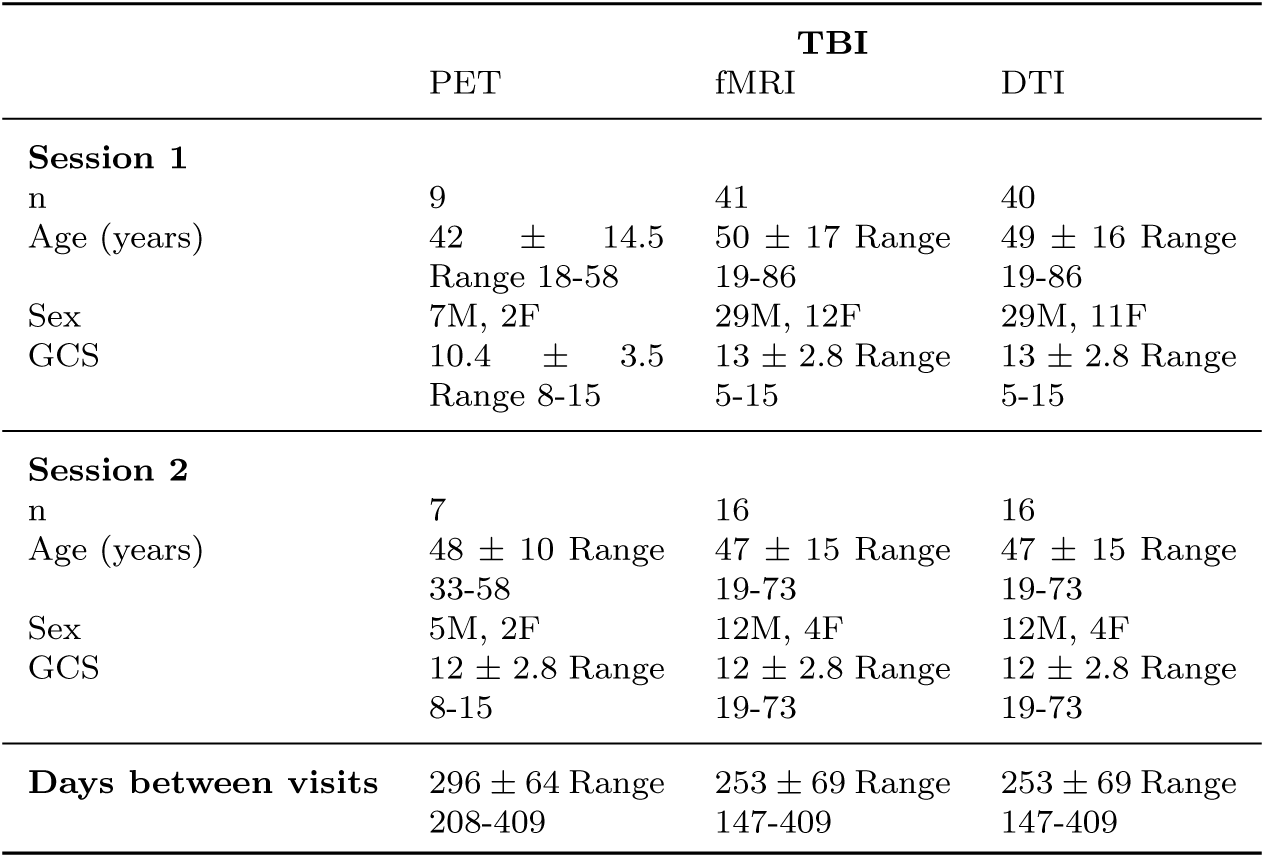
Demographic and clinical data for the TBI group.

### E.3 Further ANCOVA results

Supplementary figure 3 illustrates the *η*^2^ values for the TBI group coefficients in the ANCOVA originally shown as t-stats in Figure 2 and Figure 3. The top and bottom rows are regional *η*^2^ values for the group term (TBI vs HC) from ANCOVA conducted using all TBI subjects we had available at and, respectively (see Figure 2 in main text), while the middle row shows the *η*^2^ values for the TBI group coefficient in the ANCOVA for the subacute time point but for only those with both subacute and chronic data, as in the top row of Figure 3.

### E.4 Correlations between global imaging metrics and cognitive measures

Subjects with TBI and non-injured control participants performed the Attention Network Test (ANT),^96^ a computer-administered measure designed to examine alerting, orienting, and executive attention networks. Mean reaction time and executive attention scores were normalized and corrected for the linear effect of age based on our set of non-injured controls, including those who did not have neuroimaging data (n = 67, age 22-86, 41M/26F), and transformed to psychometric z-score such that more positive values indicate better performance. Participants also performed a battery of assessments, including: Wechsler Adult Intelligence Scalefourth Edition : Coding, Digit Span test, Letter Number Sequencing (LNS), Symbol Search, Trail Making Test A (TMT-A) and B (TMT-B), Stroop test, Rivermead post concussion questionnaire (RPQ), and Glascow Outcome Scale Extended (GOSE). Each cognitive outcome measure (except the GOSE, RPQ, and executive attention) was zscored using standardized z-scoring procedures. Each imaging metric from each modality was globally averaged across all regions and this global metric was correlated with the various cognitive outcomes via Spearman partial correlation, with sex and age as co-variates. For longitudinal analysis, global average metrics and outcome metrics were computed as a percent change, where the value at chronic was subtracted by the value at the subacute timepoint, divided by the absolute value of the sum of subacute and chronic values. Results are depicted in Supplementary Figure 4. We largely see no significant relationships except for uncorrected significance between BP*_ND_* and executive attention at the chronic timepoint, where individuals with lower BP*_ND_* had better executive attention. The change in BP*_ND_* and fALFF over time was correlated with the change in GOSE over time where more increases in BP*_ND_* and fALFF were related to better GOSE scores. At the chronic timepoint, SC is negatively correlated with TMT-A. Subject number varied by outcome measure and session, details are available in Supplementary Table 3.

**Supplementary Figure 2.**
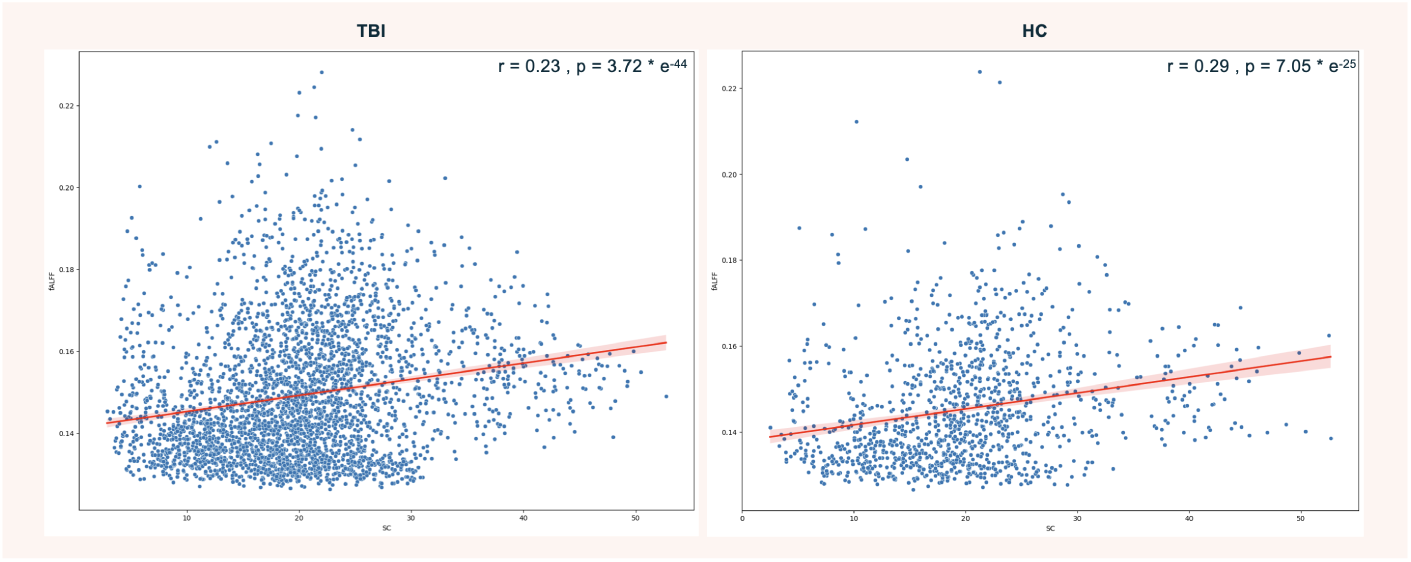
Spearman correlation between regional SC node strength and fALFF in all available healthy controls and TBI subjects at the subacute timepoint. Each point is a region from a specific individual, for all TBI subjects at the subacute time point (left) and healthy controls (right). Spearman correlation was used to examine the strength of the relationship between regional SC node strength and fALFF across all regions and individuals. Correlations are indicated in the upper right hand corner of each figure with associated permutation based p-values.

**Supplementary Figure 3.**
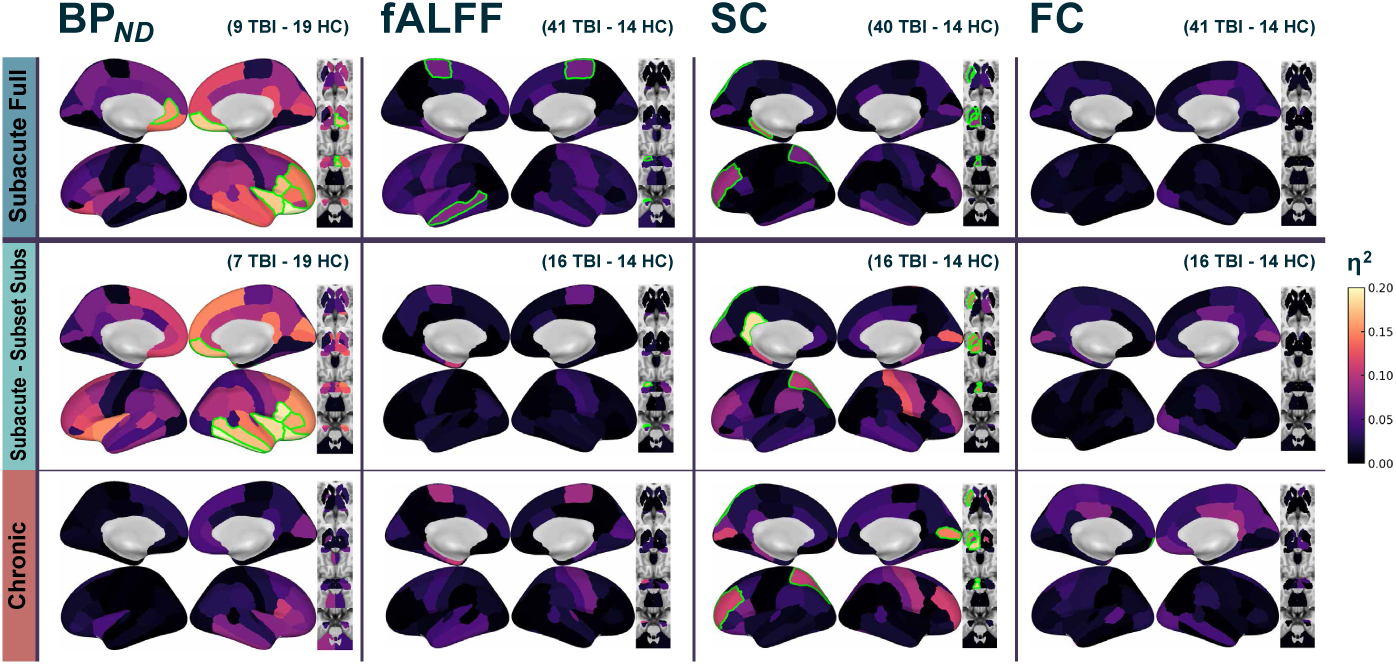
Brain plots of η values for TBI group effects from the ANCOVAs. Top row contains results for the subacute timepoint from the full group of available subjects for the subacute timepoint, while the bottom row contains results for the subjects with chronic data. The middle row are the subacute ANCOVA *η*for TBI group effects, but for only those subjects with data from both subacute and chronic time points. Regions highlighted in green reached significance at (p*<*0.05, uncorrected). N varies by modality, as indicated in each subplot.

**Supplementary Figure 4.**
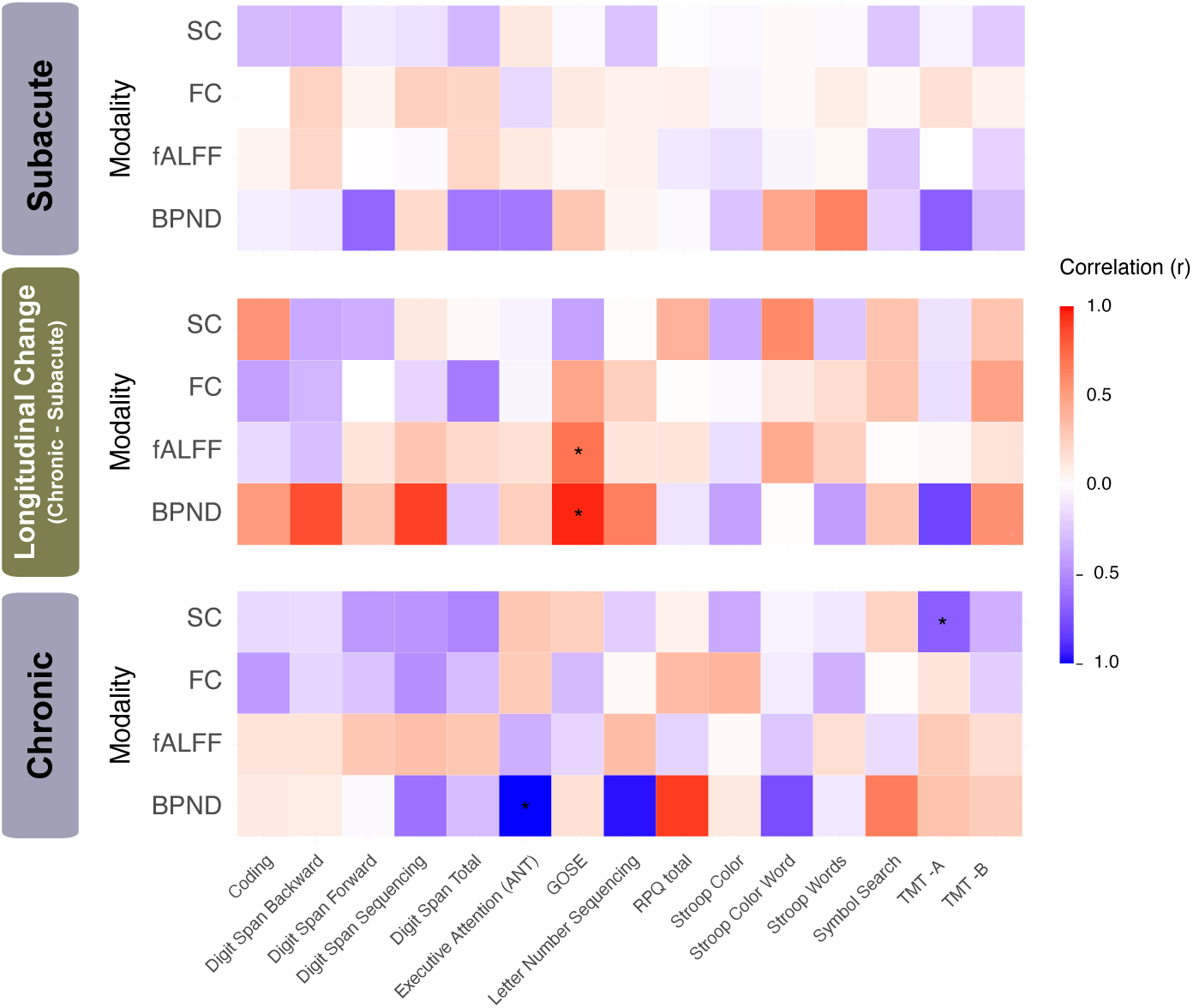
Spearman partial correlations of the wholebrain average of each modality’s metric (rows) with z-scored cognitive outcomes (columns). Spearman correlations accounted for age and sex. Global average values for every TBI subject available at both timepoints were included in the top row (Subacute) and bottom row (Chronic). Correlations of change in global metrics vs change in cognitive outcomes over time where computed using percent change for both measures. Correlations reaching uncorrected significance are indicated with an asterisk, none survived corrections.

**Supplementary Table 2.**
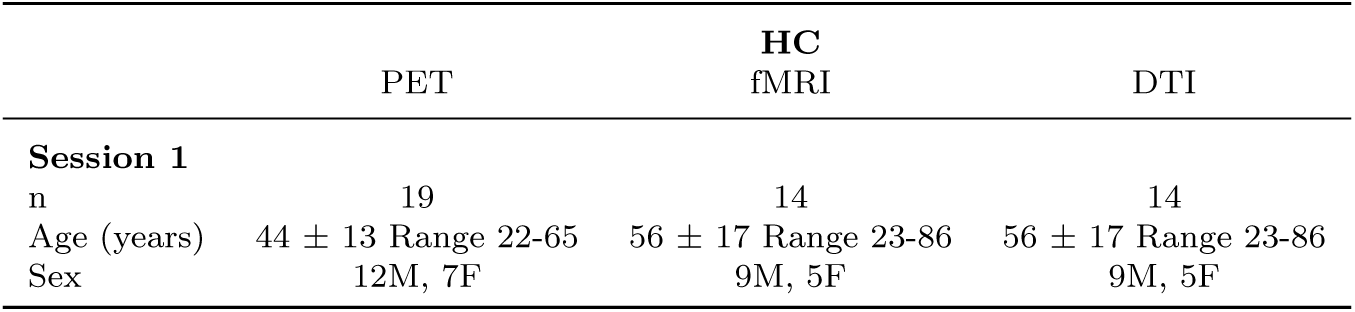
Demographic data for the HC group.

|  | PET | HC<br>fMRI | DTI |
| --- | --- | --- | --- |
| Session 1 |  |  |  |
| n | 19 | 14 | 14 |
| Age (years) | 44 ± 13 Range 22-65 | 56 ± 17 Range 23-86 | 56 ± 17 Range 23-86 |
| Sex | 12M, 7F | 9M, 5F | 9M, 5F |

**Supplementary Table 3.** Number of subjects with available data for each modality-behavior combination.

| Timepoint | Modality | Behavior or Outcome |  |  |  |  |  |  |  |  |  |  |  |  |  |  |
| --- | --- | --- | --- | --- | --- | --- | --- | --- | --- | --- | --- | --- | --- | --- | --- | --- |
|  |  | Coding | Digit Span Backward | Digit Span Forward | Digit Span Sequencing | Digit Span Total | Exec Attn | GOSE | LNS | RPQ | Stroop Color | Stroop Color Word | Stroop Words | Symbol Search | TMT-A | TMT-B |
| Session 1<br>Subacute | dMRI | 39 | 39 | 39 | 39 | 39 | 40 | 39 | 38 | 39 | 37 | 37 | 37 | 39 | 39 | 38 |
|  | fMRI | 40 | 40 | 40 | 40 | 40 | 41 | 40 | 39 | 40 | 38 | 38 | 38 | 40 | 40 | 39 |
|  | PET | 7 | 7 | 7 | 7 | 7 | 9 | 7 | 7 | 7 | 7 | 7 | 7 | 7 | 7 | 7 |
| Session 2<br>Chronic | dMRI | 13 | 14 | 14 | 13 | 14 | 16 | 13 | 11 | 15 | 13 | 13 | 13 | 13 | 13 | 13 |
|  | fMRI | 13 | 14 | 14 | 13 | 14 | 16 | 13 | 11 | 15 | 13 | 13 | 13 | 13 | 13 | 13 |
|  | PET | 7 | 7 | 7 | 6 | 7 | 7 | 7 | 6 | 7 | 7 | 7 | 7 | 7 | 7 | 7 |
| Longitudinal<br>Change | dMRI | 13 | 14 | 14 | 13 | 13 | 16 | 13 | 11 | 15 | 13 | 13 | 13 | 13 | 13 | 13 |
|  | fMRI | 13 | 14 | 14 | 12 | 13 | 16 | 13 | 11 | 15 | 13 | 13 | 13 | 13 | 13 | 13 |
|  | PET | 7 | 7 | 7 | 5 | 7 | 7 | 7 | 6 | 7 | 7 | 7 | 7 | 7 | 7 | 7 |

## Notes

### Competing Interest Statement

The authors have declared no competing interest.

